# Allele-specific binding of RNA-binding proteins reveals functional genetic variants in the RNA

**DOI:** 10.1101/396275

**Authors:** Ei-Wen Yang, Jae Hoon Bahn, Esther Yun-Hua Hsiao, Boon Xin Tan, Yiwei Sun, Ting Fu, Bo Zhou, Erc L. Van Nostrand, Gabriel A. Pratt, Peter Freese, Xintao Wei, Giovanni Quinones-Valdez, Alexander E. Urban, Brenton R. Graveley, Christopher B. Burge, Gene W. Yeo, Xinshu Xiao

## Abstract

Allele-specific protein-RNA binding is an essential aspect that may reveal functional genetic variants influencing RNA processing and gene expression phenotypes. Recently, genome-wide detection of in vivo binding sites of RNA binding proteins (RBPs) is greatly facilitated by the enhanced UV crosslinking and immunoprecipitation (eCLIP) protocol. Hundreds of eCLIP-Seq data sets were generated from HepG2 and K562 cells during the ENCODE3 phase. These data afford a valuable opportunity to examine allele-specific binding (ASB) of RBPs. To this end, we developed a new computational algorithm, called BEAPR (Binding Estimation of Allele-specific Protein-RNA interaction). In identifying statistically significant ASB sites, BEAPR takes into account UV cross-linking induced sequence propensity and technical variations between replicated experiments. Using simulated data and actual eCLIP-Seq data, we show that BEAPR largely outperforms often-used methods Chi-Squared test and Fisher’s Exact test. Importantly, BEAPR overcomes the inherent over-dispersion problem of the other methods. Complemented by experimental validations, we demonstrate that ASB events are significantly associated with genetic regulation of splicing and mRNA abundance, supporting the usage of this method to pinpoint functional genetic variants in post-transcriptional gene regulation. Many variants with ASB patterns of RBPs were found as genetic variants with cancer or other disease relevance. About 38% of ASB variants were in linkage disequilibrium with single nucleotide polymorphisms from genome-wide association studies. Overall, our results suggest that BEAPR is an effective method to reveal ASB patterns in eCLIP and can inform functional interpretation of disease-related genetic variants.

## Introduction

Facilitated by recent technological advances, numerous human genomes are being sequenced, cataloging an unprecedented amount of genetic variants (GVs)^1^. A major challenge exists in identifying and interpreting potentially functional GVs. Disease-associated GVs often reside in non-coding regions, such as introns or 3’ untranslated regions (UTRs), making it especially challenging for functional interpretation^2^. It is increasingly appreciated that many GVs in the introns or 3’ UTRs may affect RNA processing or mRNA turnover^3^. Thus, methodologies that can effectively capture functional GVs in these post-transcriptional processes are in great demand.

RNA-binding proteins (RBPs) are core players in post-transcriptional gene regulation^4, 5^. A large number of RBPs exert their function via sequence-specific protein-RNA interaction. The sequence specificity of RBPs implies that GVs may disrupt the RBP recognition of RNA substrates. Specifically, the alternative alleles of a GV may confer different binding specificity for an RBP, thus causing allele-specific functional consequences^6^. Indeed, allele-specific binding (ASB) analysis of an RBP to GVs is arguably the most direct means to identify potentially functional GVs in post-transcriptional regulation.

To detect ASB of a specific RBP, one powerful method is to examine global binding sites of the RBP. If a heterozygous GV is present within the binding site, allelic bias of the GV in the protein-bound RNA directly suggests existence of ASB. The advantage of this analysis is that the alternative alleles of a GV is examined in the same cellular environment in the same subject. Thus, the method controls for tissue conditions, trans-acting factors, global epigenetic effects, and other environmental influences.

To carry out genome-wide ASB analyses, it is necessary to capture global protein-RNA interaction in a sequence-specific manner. UV crosslinking and immunoprecipitation followed by sequencing (CLIP-Seq) is a most-often used method for this type of global profiling^7^. Recently, the enhanced CLIP (eCLIP) protocol was developed that significantly improves the efficiency and sensitivity of CLIP^8^. As a result of the improved efficiency, multiple biological replicates of eCLIP can be generated for the same experiment. In addition, each eCLIP assay is accompanied by a size-matched input (SMInput) sample as a stringent control for non-specific binding. The ENCODE project generated hundreds of eCLIP-Seq data sets for 154 RBPs in two cell lines, HepG2 and K562^9^. These data sets afford an invaluable opportunity to examine ASB patterns and shed light on the functions of GVs in post-transcriptional regulation.

However, ASB analysis is challenging in that it entails accurate quantification of single nucleotides in sequencing reads, which is easily confounded by possible inherent biases in the CLIP protocol and the limited sequencing depth available for most CLIP data sets. Nevertheless, the unique advantages of eCLIP-Seq, such as the availability of biological replicates and SMInput samples, offer an opportunity to accurately identify ASB events. Thus far, no computational method is available that leverages these unique features of eCLIP for ASB detection. Here, we present a new method called BEAPR (**B**inding **E**stimation of **A**llele-specific **P**rotein-**R**NA interaction) for this purpose. BEAPR controls for inherent bias in crosslinking using the SMInput samples, and tests for significant binding bias by taking into account the variability in the data as manifested in the biological replicates. We show that BEAPR outperforms other often-used methods for allele-specific analyses of read counts. Importantly, BEAPR is robust to overdispersion in the sequencing data and its performance is consistent across different read coverages. Applied to the ENCODE eCLIP-Seq data sets, BEAPR identified thousands of ASB events. Supported by experimental validation, these ASB events include many that can potentially cause splicing changes, alter mRNA abundance or explain the functional consequences of disease-associated GVs. Together, our results suggest that BEAPR is an effective method for ASB detection and can serve as a fundamental tool to predict functional GVs in post-transcriptional gene regulation.

## Results

### Identification of allele-specific binding of RBPs by BEAPR

BEAPR analyzes eCLIP-Seq data to identify ASB events in protein-RNA interaction. The standard eCLIP-Seq protocol generates an input control sample (SMInput) and two biological replicates of eCLIP samples^9^. As illustrated in Fig. 1a (see Methods for details), BEAPR takes as input mapped reads and peak calls from these data sets. It first identifies heterozygous single nucleotide variants (SNVs) that show bi-allelic expression in the SMInput reads. An optional input is a list of sample-specific heterozygous SNVs. If provided, this list will be combined with BEAPR-identified heterozygous SNVs. Although genome-sequencing or genotyping data may exist for the specific samples, identification of SNVs using SMInput reads may complement these data (see below).

**Figure 1.**
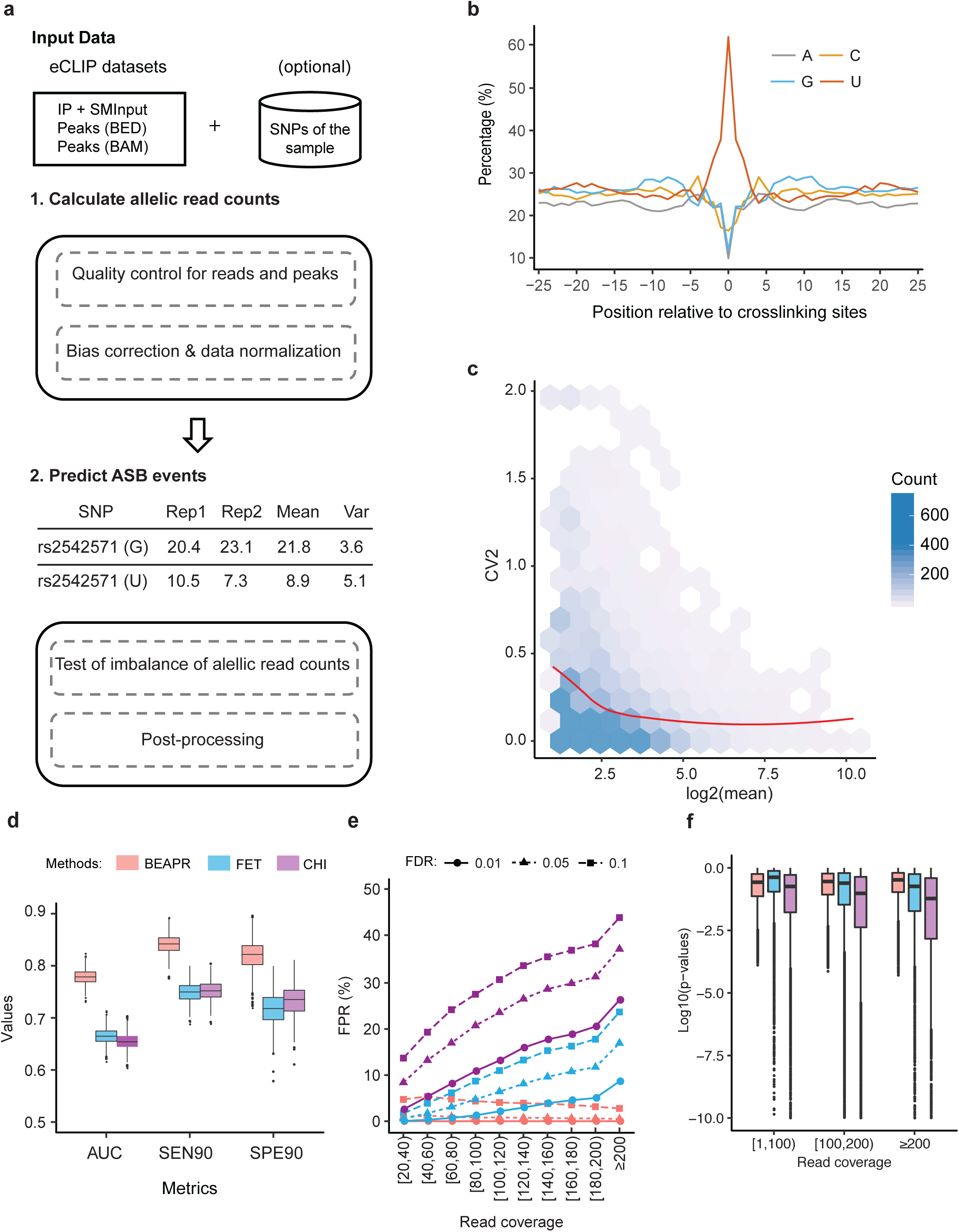
Overview of BEAPR and its performance. (a) Overall work flow of BEAPR. See Methods for details. (b) Crosslinking bias in the SMInput sample of RBFOX2 (HepG2 cells). Y axis shows the relative % of each nucleotide observed at each position relative to the crosslinking site (x = 0). (c) The square of the coefficient of variation (CV2) plotted as a function of the observed allelic read counts (mean of the two replicates) in the RBFOX2 eCLIP data in HepG2 cells (Methods). (d) Performance comparison of 3 methods using simulated data and true allelic ratio of 0.8 for ASB. Data derived from 1000 simulation experiments each encompassing 5000 SNVs. FET: Fisher’ Exact test; CHI: Chi-Squared test; AUC: area under the curve of the precision-recall curve. SEN90: sensitivity at 90% specificity; SPE90: specificity at 90% sensitivity. (e) False positive rates (FPRs) of the 3 methods using simulated data as in (d). The x-axis shows different read coverage bins. Average FPR values of all simulated data points are shown in each bin. (f) Box plots of p values calculated by the 3 methods at different levels of read coverage.

A unique feature of BEAPR is the estimation of crosslinking-induced sequence bias, using the SMInput data of each eCLIP experiment. As an example, Fig. 1b shows the bias estimation of the RBFOX2 eCLIP data generated in HepG2 cells. The enrichment of uracils at the crosslinking sites is consistent as the observations in previous CLIP studies^10^. This estimated bias, specific to each eCLIP experiment, is used to normalize the allele-specific read counts of each SNV. Subsequently, BEAPR employs an empirical Gaussian distribution to model the normalized read counts, with the expected variance estimated using a regression model (Fig. 1c, Methods). BEAPR tests whether the normalized read counts of the alternative alleles of an SNV are significantly different (i.e., existence of ASB). The predicted ASB events were subject to several post-processing filters to remove those located in homopolymeric or repetitive regions.

### Evaluation of BEAPR performance using simulated data

We first generated simulated data to evaluate the performance of BEAPR. Specifically, we carried out 1000 simulation experiments. In each experiment, 5000 heterozygous SNVs were included, each of which was assigned a total read coverage and read counts for two alternative alleles, with two simulated biological replicates. The total read coverage of the SNVs were sampled randomly from the actual read coverage distribution of SNVs of an ENCODE eCLIP data set (SRSF1 in K562) by keeping the same variance across replicates (Fig. S1). The read counts for alternative alleles of an SNV were determined using a zero-truncated negative binomial distribution given the simulated total read coverage and an expected allelic ratio *r* (read count of major allele/total read count). The value of *r* was set to be 0.5 for 90% of the simulated SNVs in each experiment. The other 10% were simulated ASB events (i.e., true positives) with an *r* value of 0.7, 0.8 or 0.9 in different experimental settings.

The performance of BEAPR was compared to those of two other methods: Chi-squared test and Fisher’s Exact test, both were used to detect allelic imbalance in read count data^11, 12^. Since these methods cannot model the variability between biological replicates, read counts from replicates were combined in using these methods. It should be noted that crosslinking-induced sequence bias was not taken into consideration by the other two methods. Thus, no such bias was simulated in our experiment, which renders some advantage to these methods. The performance of different methods was assessed by the precision-recall curves, sensitivity and specificity (Methods). As shown in Fig. 1d and Fig. S2, BEAPR achieved the highest Area-Under-the-Curve (AUC) in the precision-recall curves, and the highest sensitivity and specificity among the three methods. In general, ASB identification is challenging if the true allelic ratio is close to 0.5, given limited read coverage (Fig. S1). The performance of the three methods deteriorates at smaller *r* values (e.g., 0.7). Nevertheless, BEAPR outperformed the other methods consistently at all tested allelic ratios.

### BEAPR accounts for overdispersion in allelic read counts

Overdispersion exists if the variance of the count data is underestimated, which may lead to enrichment of very small p values and false positive predictions. In the simulation study, we examined whether the results of different methods reflected overdispersion in the data. Figure 1e and Fig. S2b&e show that the false positive rates (FPR, % false positives among all negatives) of Chi-squared test and Fisher’s Exact test were both inflated as the read coverage increased. In contrast, BEAPR demonstrated much lower FPR across all read coverage ranges, with slightly lower FPR at higher coverage. The p values resulted from the Chi-squared test and Fisher’s Exact test decreased as read coverage increased (Fig. 1f, Fig. S2c&f), which led to the higher FPR at higher read coverage ranges. These results suggest that the Chi-squared test and Fisher’s Exact test underestimate the variability in the data and thus make many false positive predictions. This overdispersion issue largely distorted the predictions made by these methods. Compared to these methods, BEAPR is robust to the variance in the input read count data, which contributed to its superior performance.

### Analysis of ENCODE eCLIP-Seq data using BEAPR

We obtained eCLIP-Seq data of 154 RBPs derived from HepG2 or K562 cells as part of the ENCODE project^9^. Each RBP had two biological replicates of eCLIP and one SMInput control. The reads were pre-processed and mapped using STAR as described previously^8^. eCLIP peaks were identified using CLIPper^13^. In this work, eCLIP peaks were retained for subsequent analyses if the read coverage in at least one replicate is ≥4-fold of that in the corresponding region in the SMInput.

Given the above defined eCLIP peak regions, BEAPR proceeds to identify heterozygous SNVs in these regions, which will be combined with sample-specific SNVs if provided by the user. For the ENCODE data sets, we identified heterozygous SNVs in HepG2 and K562 cells using both the eCLIP data and whole-genome sequencing data^14^ (Methods). Within the eCLIP peak regions, the heterozygous SNVs identified via the two methods overlapped substantially (Fig. 2a). Around 93.1% of eCLIP-derived SNVs in HepG2 (89.5% in K562) were also identified in the respective whole-genome sequencing data. About 80.4% of SNVs located in eCLIP peaks and predicted by whole-genome sequencing of HepG2 (71.9% for K562) were also identified by our method. Thus, if assuming whole-genome sequencing as the ground truth, eCLIP-based SNV identification achieved a precision of 93.1% and 89.5%, and a sensitivity of 80.4% and 71.9%, in HepG2 and K562 for SNVs located in eCLIP peaks, respectively. Furthermore, we experimentally confirmed 5 heterozygous SNVs that were identified in the eCLIP data, but missed by whole-genome sequencing (Fig. S3), supporting the validity of these SNVs. Our results suggest that BEAPR can effectively identify heterozygous SNVs using eCLIP data alone, if genotyping or genome sequencing data are not available for the specific sample.

**Figure 2.**
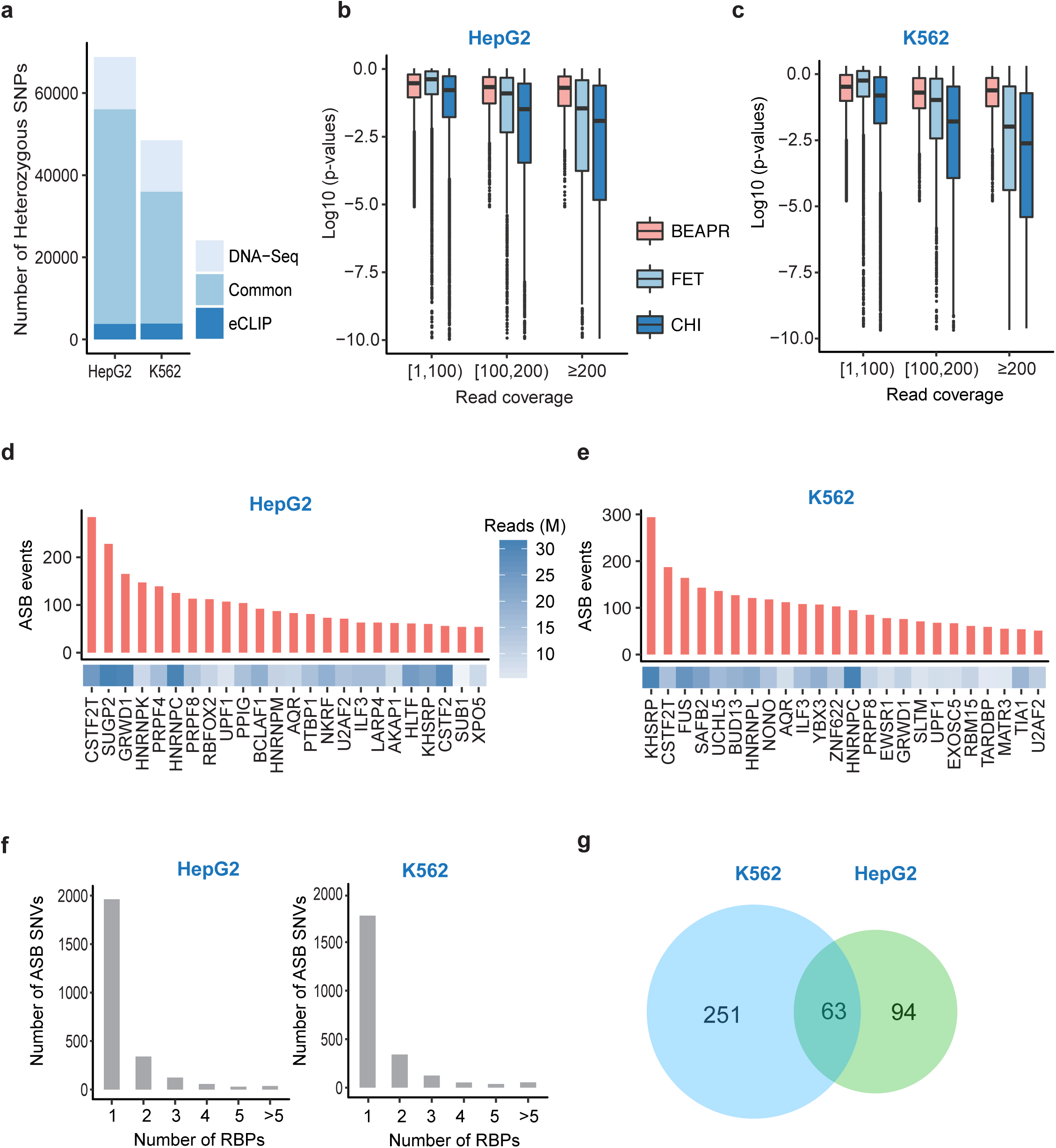
ASB events identified in eCLIP data of HepG2 and K562 cells. (a) Number of heterozygous SNVs identified via whole-genome DNA sequencing (DNA-Seq) or eCLIP. Those that are common to both methods are illustrated. (b) P values calculated by the 3 methods at different levels of read coverage in HepG2. (c) Same as (b), for K562 cells. (d) Number of ASB events identified for each RBP in HepG2. Only RBPs with ≥50 ASB events are shown. The number of usable eCLIP-Seq reads (in millions (M)) is shown for each RBP. (e) Same as (d), for K562 cells. (f) Number of ASB SNVs associated with 1 or more than 1 RBPs. (g) The overlap of ASB SNVs between HepG2 and K562. Only heterozygous SNVs common to the two cell lines are included.

Next, we asked whether the prediction of ASB by BEAPR reflected overdispersion in the ENCODE eCLIP data. As shown in Fig. 2b and c, BEAPR yielded relatively stable p values at different levels of read coverage. Similar to the observations in simulated data, Chi-squared test and Fisher’s Exact test suffered from overdispersion especially at higher levels of read coverage. The results suggest that these methods underestimate the variance of allelic read counts in the eCLIP data and tend to produce false positive predictions given high read coverage.

### ASB events identified in ENCODE eCLIP-Seq data

For all RBPs with eCLIP data, a total of 3706 and 3783 ASB events were identified in the HepG2 and K562 cells, respectively. The RBPs with more than 50 predicted ASB events are illustrated in Fig. 2d and e and the results for all RBPs are shown in Fig. S4. The numbers of ASB events associated with different RBPs varied greatly. This variation may be accounted for by multiple factors, such as sequencing depth, number of eCLIP peaks and the binding specificity of the RBP. The genomic distribution of ASB events often reflects known functions of the RBPs (Fig. S5). For example, proteins known to regulate RNA stability, such as UPF1^15^, showed ASB enrichment in the 3’UTR regions. ASB events of known splicing regulators^15^, such as RBFOX2, PTBP1, PRPF8, U2AF1 and heterogeneous nuclear ribonucleoproteins (hnRNPs), were enriched in the intronic regions.

The above ASB events occurred in 2552 and 2384 SNVs in HepG2 and K562, respectively. As shown in Fig. 2f, 76.9% and 77.6% of these SNVs were associated with ASB of one RBP in HepG2 and K562, respectively, suggesting that ASB events are mostly RBP-specific. Nevertheless, multiple RBPs may interact leading to common eCLIP peaks and ASB events.

Next, we compared the ASB events between the two cell lines. A total of 762 heterozygous SNVs were testable for ASB in both cell lines. Among these SNVs, 157 and 314 were identified with ASB patterns in HepG2 and K562, respectively, with 63 shared by the two cell lines (Fig. 2g, p = 9.7e-27). This result supports that GVs affecting protein-RNA binding could function in a cell-type independent manner, at least between the two cell lines tested in this study.

### SNVs with ASB patterns disrupt RBP binding motifs

For an RBP with specific sequence preference, it is expected that ASB patterns may arise if an SNV disrupts its binding motif. Thus, the localization of ASB SNVs near known RBP binding motifs serves as a strong indicator of the validity of the predicted ASB. We obtained binding motifs identified by the RNA Bind-n-Seq (RBNS) assay as part of the ENCODE project^9^. RBNS quantifies the binding specificity of an RBP to a k-mer sequence using the R value^16, 17^. To focus on RBPs with relatively high binding specificity, we required an RBP to have at least one pentamer sequence with R ≥ 2. Among all RBPs with at least 50 ASB events and at least 30 events in annotated genes, five RBPs, hnRNPC, hnRNPK, hnRNPL, RBFOX2 and TARDBP had RBNS pentamers that passed this requirement. For these RBPs, we analyzed the occurrence of the RBNS pentamers in the flanking regions of ASB SNVs. Compared to control regions (Methods), the ASB flanking regions were enriched with RBNS pentamers, and importantly, the ASB loci were located in close proximity to the pentamer enrichment peaks (Fig. 3a-f). Interestingly, the fold enrichment of RBNS pentamers in ASB regions relative to control regions is higher for proteins with higher R values based on RBNS. These results strongly support the validity of the predicted ASB events. We observed that the specific nucleotide position disrupted by the ASB SNVs varied for different RBPs (Fig. 3a-f), which may depend on the specific binding property of the RBPs and the specific allelic sequences of ASB SNVs.

**Figure 3.**
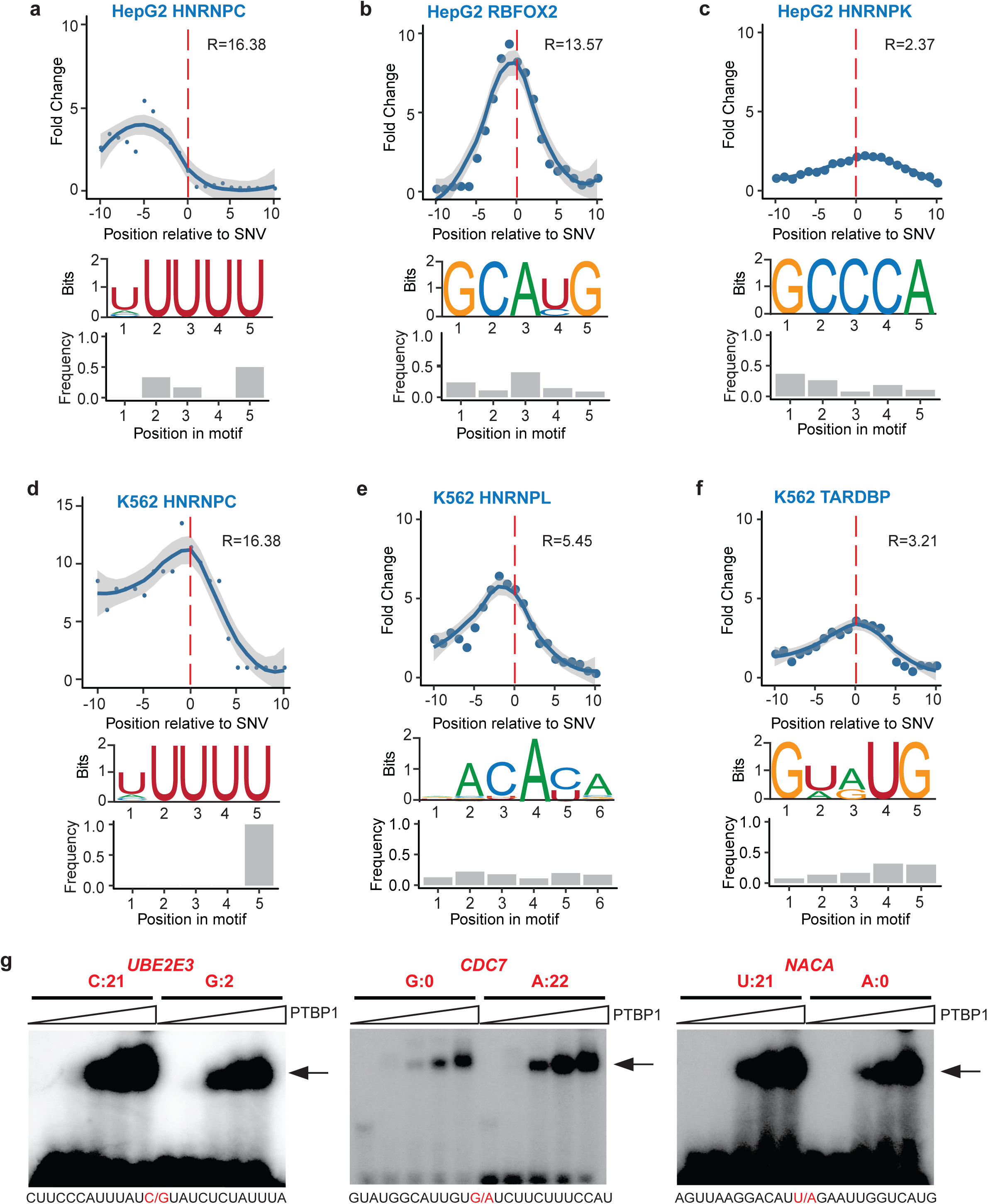
Bioinformatic and experimental validation of ASB. (a-f) Enrichment of RBNS motifs in the regions around ASB SNVs (x = 0) of each RBP. Y axis shows fold change in the enrichment relative to randomly chosen control regions (Methods). Ten sets of controls were constructed, with the regression curve and 95% confidence interval of the average fold change shown in the plot. The highest R value among the RBNS pentamers for each RBP is shown. The RBNS pentamer sequences are visualized in the pictograms. The relative frequency of the ASB SNVs overlapping each motif position is shown in the bar graph. (g) EMSA results of PTBP1 binding to its ASB targets. Alternative alleles of the ASB SNVs were synthesized, as labeled above the gel images. The read counts of the alternative alleles are shown. The sequences of the synthetic RNA fragments are shown below each gel image, where the ASB SNV is highlighted in red. The arrow indicates RNA-protein complex. Increasing concentrations of PTBP1 were used in different lanes of the gel image (from left to right: 0, 0.6, 1.2, 2.5, and 5 μg).

For RBPs with ≥50 ASB events but without specific RBNS motifs, we identified the top five most-frequent pentamers in the 21mer region centered at the ASB SNV of each RBP (Fig. S6). These pentamers are largely consistent with general binding preferences of RBPs known in the literature, such as those for hnRNP L (CA-rich motif^18^) and PTBP1 (CU-rich motif^19^). In addition, the positional distribution of these enriched pentamers often showed biases relative to the loci of ASB SNVs. This bias is evident for proteins with well-known sequence specificity in their binding preference (such as PTBP1). As expected, proteins with low binding specificity may not demonstrate strong signals in this analysis. These results again support that ASB analysis can effectively capture specific RBP binding sites and allelic biases in protein-RNA interaction.

### Experimental validation of ASB events

To provide direct experimental support that ASB SNVs alter the binding of RBPs, we carried out electrophoretic mobility shift assays (EMSA, or gel shift) on randomly selected ASB events of PTBP1 (Fig. 3g, Fig. S7). This protein was chosen since it is relatively easy to purify. To confirm that the ASB SNVs alter the binding of PTBP1, two versions of each target RNA were synthesized harboring the alternative alleles of the SNV. As shown in Fig. 3g, the binding of PTBP1 to target RNAs was stronger with increasing protein input. Strong signals of differential binding to the alternative alleles of the SNVs were observed for all three RNA targets. In addition, the alleles with the stronger gel shift signals corresponded to the alleles with more eCLIP reads, supporting the validity of the predicted ASB events.

Together, the above results support the validity of our ASB identification method. Since ASB serves as a direct indicator of functional SNVs, it is expected that ASB patterns can inform functional interpretations of GVs. Next, we examined whether the above ASB analysis captured functional SNVs in regulating alternative splicing and mRNA abundance.

### SNVs subject to ASB by splicing factors may cause splicing alteration

The functional consequence of the ASB event depends on the function of the RBP. Since many RBPs in this study are known splicing factors, we examined whether some ASB events may alter splicing. We collected all ASB events of known splicing factors in each cell line (37 in HepG2 and 31 in K562). First, we examined the distance of intronic SNVs with ASB by splicing factors to the nearest splice site. Compared to randomly selected SNVs in the same introns (Methods), ASB SNVs were significantly closer to the splice sites (Fig. 4a). Next, to verify that the ASB events are associated with regulatory targets of the splicing factors, we analyzed the splicing changes of the associated exons upon knockdown (KD) of the corresponding RBP using ENCODE RNA-Seq data in HepG2 or K562 cells^9^. Compared to random controls (Methods), ASB-associated exons had a significantly larger change in the percent spliced-in (PSI) values upon KD of the splicing factors (Fig. 4b). This result supports that ASB-associated exons are bona fide targets of the splicing factors. It should be noted that PSI changes of the ASB target exons upon splicing factor KD are not expected to be very large in magnitude because the nature of ASB implicates that only one of two alleles of the endogenous SNV is bound strongly by the corresponding RBP.

**Figure 4.**
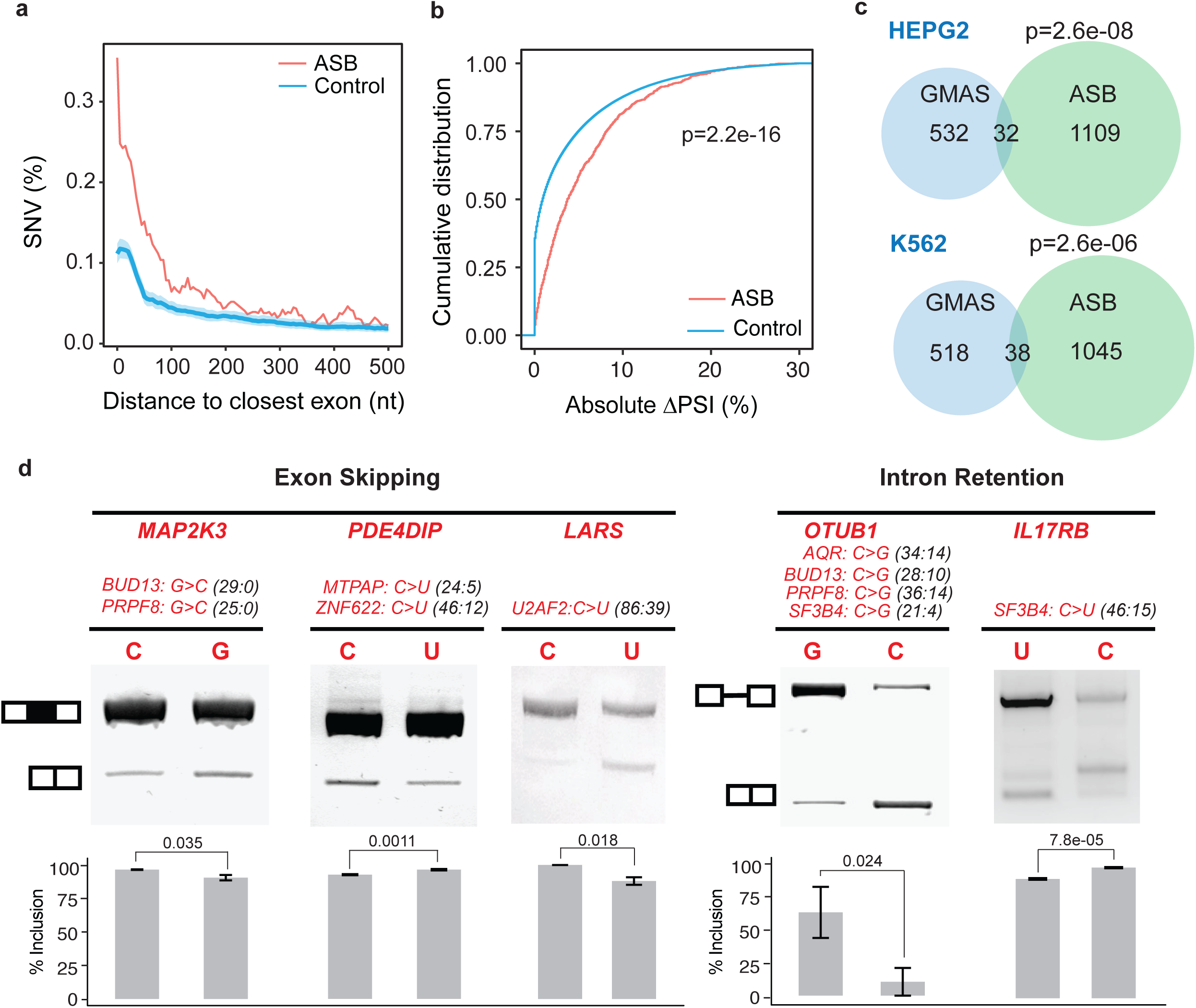
Function relevance of ASB SNVs in splicing regulation. (a) Distance of intronic ASB SNVs to the nearest splice sites. Controls are randomly chosen SNVs in the same introns. 100 sets of controls were constructed, with the average and standard deviation shown in the plot. (b) Absolute change in the PSI values of exons associated with ASB events of splicing factors upon knockdown of the respective splicing factor in HepG2 or K562 cells. Controls were chosen as random intronic SNVs in the same introns as the tested ASB SNVs. P value was calculated by the Kolmogorov-Smirnov test. (c) Overlap of ASB SNVs of splicing factors with heterozygous SNVs associated with GMAS events in the genes harboring ASB SNVs. GMAS-related SNVs were required to reside in the GMAS exon or within 500nt from exon-intron boundaries. P values were calculated via the hypergeometric test, with the background as the total number of heterozygous SNVs in genes with ASB SNVs. (d) Splicing reporter validation of the function of ASB events. Three exon skipping and two intron retention events are included. The gene names (bold) with the ASB events (alleles and read counts) and the associated RBPs are shown. Inclusion level (mean+/-SD, 3 biological replicates) of the exon or intron is shown below each gel image. P values were calculated by Student’s *t*-test.

### ASB-associated SNVs have significant association with allele-specific splicing events

If an ASB event is functional, we expect that the associated exon or gene is under cis-regulation by this SNV. For splicing factors, such ASB events will lead to allele-specific alternative splicing (i.e., genetically modulated alternative splicing, GMAS). Using our previous methods^6, 20^, we identified GMAS events in RNA-Seq data of control HepG2 and K562 cells generated by the ENCODE project. Using these data, we observed that SNVs with ASB patterns are significantly enriched in the GMAS exons or within 500nt from their exon-intron boundaries (Fig. 4c). This observation supports the hypothesis that splicing factor-associated ASB imposes cis-regulation to splicing.

### Experimental validation of ASB-associated SNVs for splicing regulation

Based on the above results, it is very likely that ASB SNVs of splicing factors are causal GVs responsible for genetic regulation of alternative splicing. To provide experimental support for this hypothesis, we tested five ASB SNVs regarding their potential impact on alternative splicing. These events were chosen to include ASB SNVs located in exons or within 500nt away from exons. For each ASB SNV, the relevant exonic and intronic regions were cloned into a minigene reporter^6^ (Methods). Two minigenes were created for each SNV, harboring the two alternative alleles respectively. Upon transfection into HeLa cells, splicing of the middle exon was analyzed using RT-PCR with primers targeting the two GFP exons (Fig. 4d). All five exons were confirmed to have allele-specific splicing patterns. Note that the direction of splicing enhancement or repression by each allele depends on the specific associated RBP.

### SNVs subject to ASB in 3’ UTRs may regulate RNA abundance

Many RBPs regulate RNA abundance by binding to cis-regulatory elements in 3’ UTRs^21^. Among all RBPs with ASB events in 3’ UTRs, UPF1 had the highest number of events in both cell lines (Fig. S8a). Given UPF1’s well-known function in RNA degradation^15^, we asked whether genes with these ASB events demonstrated expression changes upon UPF1 KD. Using ENCODE RNA-Seq data sets, we analyzed differential expression of genes with UPF1 ASB in their 3’ UTRs. Figure 5a shows the false discovery rate (FDR) values of the differential expression test of target genes and random controls. The results support that UPF1 regulates gene expression via ASB to SNVs. Note that the magnitude of gene expression change is not large given the heterozygotic nature of the SNVs in the cells. Nevertheless, the gene expression changes of UPF1-ASB targets are significantly larger than those of control genes (Fig. S8).

**Figure 5.**
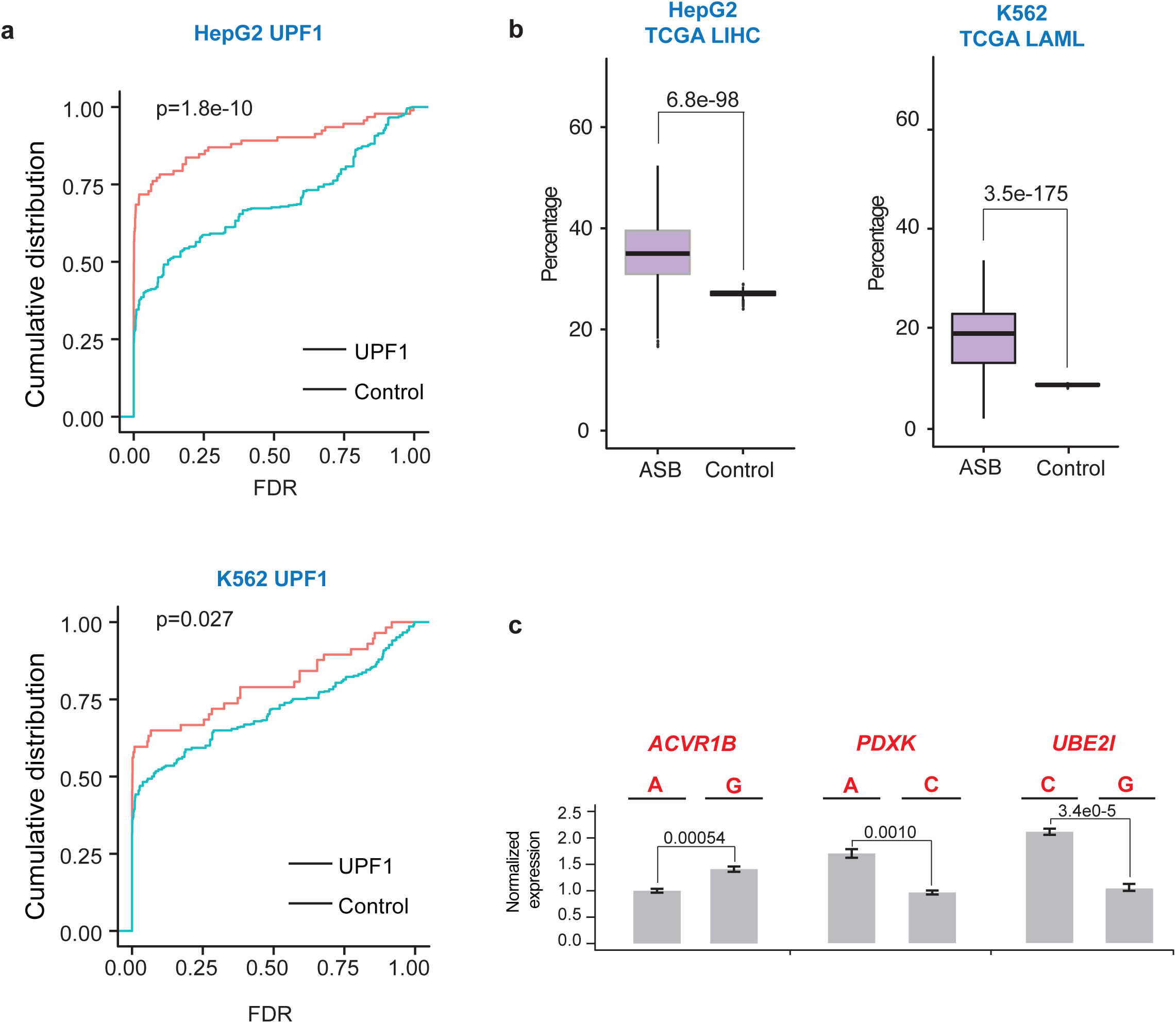
Functional relevance of ASB SNVs in regulating mRNA abundance. (a) Distributions of FDR values of the differential gene expression upon UPF1 knockdown in HepG2 or K562 cells. Data for genes with ASB SNVs of UPF1 in their 3’ UTRs and control genes are shown, where the controls were chosen as genes without UPF1 eCLIP peaks and with similar expression levels as UPF1 targets (within +/-30% of RPKM). P values were calculated by the Kolmogorov-Smirnov test. (b) Percentages of eQTL genes among those genes with ASB patterns of any RBPs in HepG2 or K562. eQTL genes were extracted from the TCGA project for Liver hepatocellular carcinoma (LIHC) and Acute Myeloid Leukemia (LAML), respectively, to match the cell type of HepG2 and K562. Control is shown for the overall fraction of eQTL genes in each cancer type. P values were calculated by pair-wise *t*-test. (c) Expression of minigenes carrying alternative alleles of ASB SNVs in the 3’ UTR of mCherry. mCherry expression (mean+/-SD) measured via real-time qRT-PCR was normalized by that of eYFP (driven by bi-directional promoters). Three biological replicates were analyzed. P values were calculated by Student’s *t*-test.

### ASB-associated SNVs are significantly enriched in genes with eQTL patterns

To further examine the association of ASB with cis-regulation of gene expression, we analyzed the enrichment of ASB-associated SNVs in genes with known eQTL patterns. To this end, we obtained eQTL and genotype data of 792 and 400 individuals with Acute Myeloid Leukemia (LAML) and Liver Hepatocellular Carcinoma (LIHC), respectively, generated by the TCGA^22^ project. Next, we asked whether genes harboring ASB SNVs are more likely eQTL genes than expected by chance. As shown in Fig. 5b, the fraction of eQTL genes among those with ASB patterns is much higher than the overall fraction of eQTL genes in TCGA. It should be noted that since ASB patterns were identified in HepG2 and K562 cells, the observed overlap with eQTL genes here may be an underestimation of the actual overlap, given possible existence of cell-type specificity for certain ASB events. Therefore, the results here strongly support that ASB of RBPs likely impose regulation on mRNA abundance.

### Experimental validation of ASB-associated SNVs in the regulation of RNA abundance

To experimentally test the roles of ASB SNVs in regulating RNA abundance, we carried out reporter assays for 3 events randomly chosen from all ASB SNVs located in 3’ UTRs. The reporter has a bi-directional promoter that drives the expression of mCherry and eYFP (Methods). Regions of the 3’ UTRs flanking the ASB SNVs were cloned as the 3’ UTR of mCherry (Methods), while eYFP serves as an internal control for gene expression. For each SNV, two versions of the reporter were constructed carrying each of the two alternative alleles. Upon transfection to HeLa cells, we measured mCherry and eYFP expression via real time qRT-PCR. As shown in Fig. 5c, all 3 SNVs were confirmed as causal differential RNA abundance of the reporter. These results support that ASB analysis could help to pinpoint functional varaints in regulating gene expression.

### SNVs with ASB patterns overlap disease-associated genetic variants

To examine whether ASB by RBPs may explain the functional roles of disease-related genetic variants, we compared ASB SNVs in this study with known disease-associated GVs included in several databases: GWAS, COSMIC, ClinVar, CIVIC and iGAP. As shown in Fig. 6a, a total of 89 unique ASB SNVs have known disease relevance according to this analysis. In addition, 1854 ASB SNVs (38.4% of the 4825 ASB SNVs in total combining data from HepG2 and K562) were in linkage disequilibrium (LD) with 1415 GWAS single nucleotide polymorphisms (SNPs) (and within 200kb in distance) (Fig. 6b). This high percentage of ASB SNVs with GWAS association supports the potential functional relevance of ASB. Among these GWAS SNPs, 29% were in the same genes as the ASB SNVs (Fig. 6b). The vast majority of the GWAS SNPs were located in introns whose functional consequence had been hard to predict.

**Figure 6.**
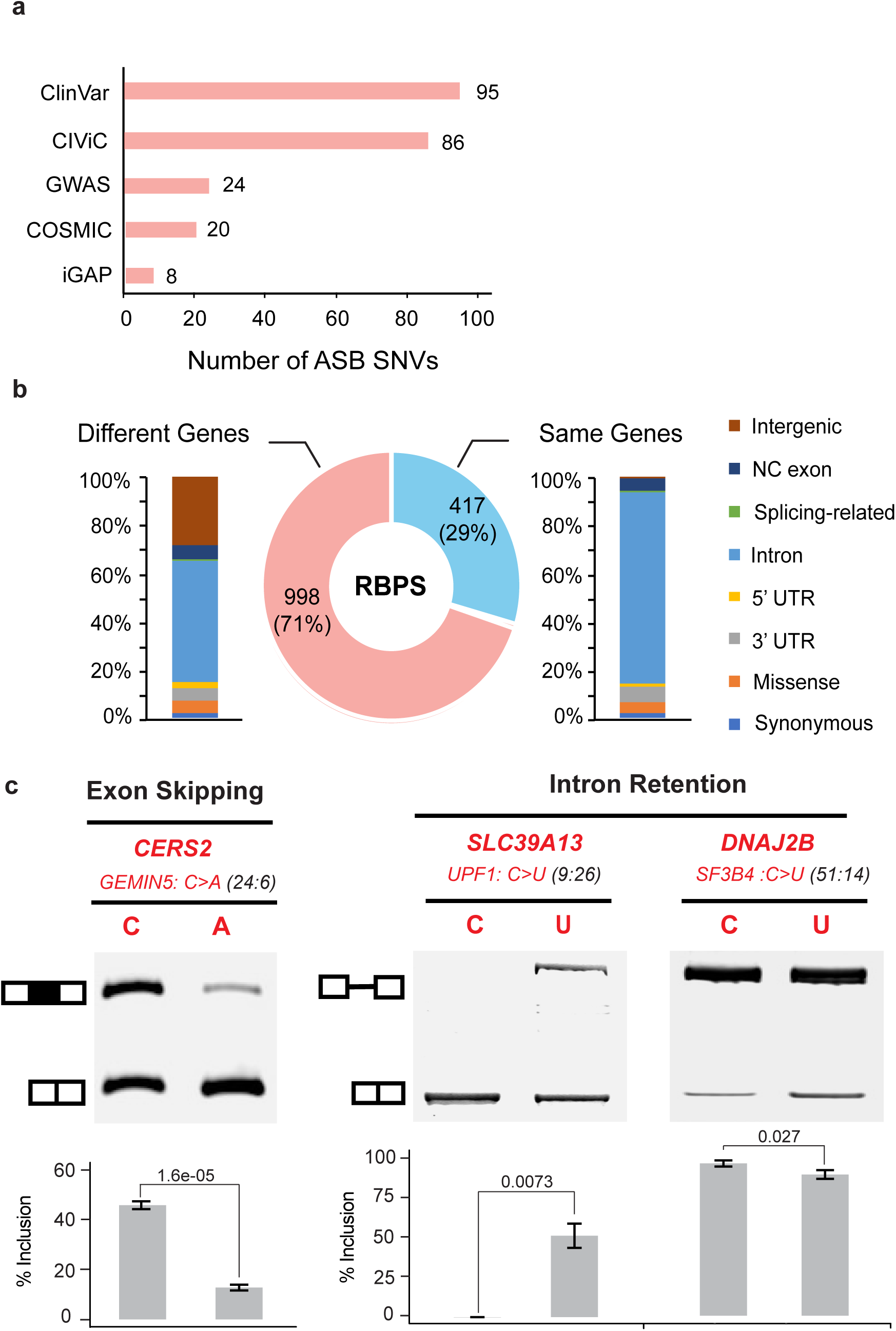
ASB events inform functional interpretation of disease-associated variants. (a) Numbers of ASB SNVs that are also disease-related SNVs annotated by different databases. (b) Genomic context of GWAS SNPs (stacked bars) located in the same or different genes as ASB SNVs. Most GWAS SNPs are located in introns, whose function was elusive. NC exon: exons in non-coding transcripts. Splicing-related: those located in splice site signals. (c) Splicing reporter validation of 3 ASB SNVs, similar as Fig. 4d.

Since many disease-associated ASB SNVs are located in introns, we experimentally tested 3 events among those that are associated with splicing factors for their impact on alternative splicing. These events were picked randomly but requiring the SNVs to reside in exons or within 500nt to exon-intron boundaries. The splicing reporters were constructed similarly as described above. As shown in Fig. 6c, all 3 SNVs were confirmed as splicing-altering variants. Specifically, the SNP rs3731896 in the gene DNAJ2B was identified as an ASB SNV of the RBP SF3B4 (also called SAP49). It is a known GWAS SNP associated with educational attainment^23^. SF3B4 encodes for a subunit of the splicing factor SF3B^24^ that is part of the spliceosomal complex. In the reporter assays, we observed that this SNP alters intron retention of its host intron. The SNP rs267738 is associated with multiple traits in GWAS, such as blood protein level and rhegmatogenous retinal detachment^25, 26^. It demonstrated ASB pattern in the eCLIP data of GEMIN5, a small nuclear RNA (snRNA)-binding component of the survival of motor neurons (SMN) complex^27^. This SNP is located in the gene CERS2 with known functional relevance in cancer^28, 29^. Interestingly, the SNP was annotated as a missense variant in the GWAS catalog. Our data demonstrated significant splicing changes caused by this SNP, suggesting that exonic SNVs that appear to be nonsynonymous could function by altering splicing, an aspect that has been largely overlooked. The SNP rs2293577 is located in the gene SLC39A13 that is annotated to be associated with Alzheimer’s Disease by the IGAP consortium^30^. Intriguingly, this SNP, located in an alternatively spliced intron of the 3’UTR of SLC39A13, caused significant splicing change.

## Discussion

In this study, we developed a new method called BEAPR to identify ASB events using eCLIP-Seq data. eCLIP-Seq captures transcriptome-wide protein-RNA interaction profiles. Compared to previous CLIP methods, eCLIP improves the efficiency and reproducibility in library generation and yields high usable read percentages across diverse RBPs^8^. The large number of eCLIP-Seq data sets made available by the ENCODE project, with biological replicates and paired size-matched input controls, affords a unique opportunity to examine protein-RNA interaction in an allele-specific manner.

Quantitative analysis of SNVs in CLIP reads is challenging in that the read coverage of a single nucleotide is relatively low compared to that in an average RNA-Seq data set. In addition, technical biases, such as those due to crosslinking, may confound the estimated allelic bias and lead to false positive ASB predictions. BEAPR addresses these potential issues by modeling read count variability and crosslinking bias and filtering for other possible technical biases. Using simulated reads, we demonstrated that BEAPR out-performs standard methods for read count comparisons. Importantly, we observed that the other methods suffered from overdispersion in both simulated and actual eCLIP data, which led to their high false positive rates. In contrast, BEAPR is robust to the variance in the input read count data and overcomes the overdispersion issue.

BEAPR identified thousands of ASB events using the ENCODE eCLIP data. Supported by experimental validations, we demonstrated that the ASB patterns helped to inform functional predictions of SNVs in regulating alternative splicing and mRNA abundance. The majority of these SNVs are located in the introns or 3’ UTRs. Thus, these results helped to address one of the most challenging questions in the post-genomic era: the functional relevance of non-coding variants in human genomes.

It is increasingly appreciated that non-coding variants may affect post-transcriptional gene regulation and many contribute to disease-related processes^3^. For example, it was estimated that 35% of disease-causing point mutations disrupt splicing by altering splice site signals or auxiliary regulatory elements in the exons or introns^3^. Despite the widely recognized importance, systematic prediction of causal GVs that alter post-transcriptional gene regulation has been a major challenge. Most methods, such as splicing or gene expression QTL analyses, rely on detection of correlative relationships without the capacity to pinpoint the exact causal GVs. ASB of RBPs provides a direct means to interpret the function of SNVs. The ASB-based analyses presented in this study demonstrated that it is highly feasible to employ this approach for causal SNV detection in post-transcriptional regulation.

In addition to splicing and mRNA abundance, ASB patterns may help to identify functional SNVs involved in other aspects of RNA metabolism, such as RNA localization or RNA secondary structures. The specific function of SNVs should depend on the roles of the RBPs demonstrating ASB patterns. Therefore, we expect that allele-specific analyses of eCLIP will be an essential approach to deciphering the function of non-coding variants in the RNA.

## Methods

### Preprocessing of the ENCODE eCLIP data

eCLIP data sets generated from the HepG2 and K562 cell lines were downloaded from the ENCODE data portal. Raw reads were demultiplexed, adapter-trimmed and mapped according to established eCLIP data processing procedures of the ENCODE project^9^. After removal of PCR duplicates, the remaining uniquely mapped reads were called ‘usable’ reads for ASB analysis. eCLIP peaks were identified using read 2 (*R*_2_) of the paired-end reads via CLIPper^13^, with options -s hg19 -o -bonferroni -superlocal-threshold-method binomial-save-pickle. In this work, eCLIP peaks were retained for subsequent analyses if the library-normalized read coverage in at least one replicate is ≥4-fold of that in the corresponding region in the SMInput.

### Identification of crosslinking sites in eCLIP peaks

To examine the potential existence of crosslinking bias, BEAPR first determines crosslinking sites within eCLIP peaks. A crosslinking site is expected to coincide with the start position of a significant number of reads compared to random expectations. To identify crosslinking sites, for each eCLIP peak, the 5’ end positions of usable *R*_2_ reads within the peak were identified. For each nucleotide position *i* in the peak, the number of *R*_2_ reads, *m_i_*, whose 5’ end coincides with position *i* was obtained. The actual *m_i_* values were compared to random expectations obtained by permuting the positions of eCLIP reads within a peak. Empirical FDR was calculated and a minimum FDR of 0.001 was used to call crosslinking sites.

### Normalization of allele-specific read counts by crosslinking bias

To identify inherent crosslinking bias for an eCLIP experiment, BEAPR calculates the relative abundance of the four nucleotides at each position flanking the crosslinking sites in the SMInput sample (Fig. 1b). The observed relative sequence bias in the SMInput is unlikely resulted from the binding preference of specific RBPs. This bias is thus referred to as the crosslinking bias, which is estimated for each experiment. It is used to normalize the observed allele-specific read counts of SNVs in the vicinity of crosslinking sites.

Prior to read count normalization for each SNV, we removed low-quality read bases by requiring a minimum Base Alignment Quality (BAQ) score of 10^31^. After this procedure, we examined the 51-nt region flanking the crosslinking site in each eCLIP peak. For each offset position *d*, ‒25 ≤ *d* ≤ 25, relative to the crosslinking site, the crosslinking bias *q*(*a, d*) of the nucleotide *a* at the offset position *d* was calculated as described above. Let *y_i,a,j_* be the number of eCLIP reads mapped to the allele *a* at the SNV *i* in the eCLIP replicate *j*. *R_2_* denotes the total number (in millions) of eCLIP reads in the replicate. The normalize read count *x_i,a,j_* of *y_i,a,j_* was calculated as *x_i,a,j_* = *y_i,a,j_*/(*q*(*a, d*)×*R_j_*). If *d* > 25 or *d* < ‒25, *q*(*a, d*) was set to be 0.25.

### Estimation of expected variance of normalized read counts

For each allele *r* at a heterozygous SNV *i*, let the variance of the normalized read counts across the eCLIP replicates be 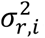 Since only a small number of replicates are available, the sample variance 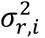 is a poor estimator of the expected variance of the normalized read counts for the allele *r*. Hence, we developed the following procedure that considers allelic read counts from all SNVs to enhance the estimation of the expected variance. Specifically, for each allele *r* at an SNV *i* located in an eCLIP peak, we calculated the mean *μ_r_,_i_* and variance 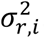 of its normalized read counts across the CLIP replicates. Using each pair of the mean and variance values, *μ_r,i_* and 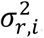 we calculated the square of the coefficient of variation (CV2) value, *ω_r,i_* Then, a LOESS regression function was applied to fit all CV2 and log2*μ_r,i_* values, where the CV2 was the response variable and the log2-scaled mean was the explanatory variable (Fig. 1c). To predict the expected variance of the normalized read counts for an allele *r*′, let 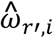 be the CV2 value moderated by the LOESS regression function. The expected variance 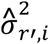 for the allele *r*′ at an SNV *i* was calculated as.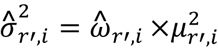

### Identification of ASB events

Let *X_A,i_* = {*x*_*A,i*,1,_… *x*_*A,i,k*_} be the normalized read counts for an allele *A* at an SNV *i* in all the *k* CLIP replicates. Similar to a previous study^32^, we used Gaussian distribution, which can adapt to different types of read count data with high or low dispersion heterogeneity, to model the normalized read counts. That is, we assume

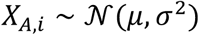

where μ is the mean and σ^2^ the variance of the Gaussian distribution. In addition, we assume the prior distribution of μ follows a normal distribution:

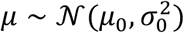

where the mean *μ*_0_ and the variance 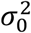 are hyper-parameters.

Let *r* and *a* denote the reference and alternative allele at an SNV site *i*. Our null hypothesis is that *r* and *a* are equivalently represented in the associated eCLIP peak. To test the null hypothesis, we examined whether *μ_r,i_* = *μ_a,i_* given *X_r,i_* and *X_a,i_*. An alternative method to test the difference of the two mean values is *t*-test. However, *t*-test was shown to be inapplicable to genomic read counts data derived from a limit number of replicates^32^. Hence, we derived the following empirical Gaussian distribution to test the null hypothesis. At an SNV *i*, an allele was named the major allele, denoted as *M*, if its average allelic read counts across the CLIP replicates was higher than that of the other allele. Otherwise, the allele was called the minor allele, denoted as *m*. In BEAPR, we calculate the empirical probability *P*(*μ* = *μ_m,i_*|*X* = *X_M,i_*) that the average read count of the minor allele, *μ_m,i_*, was generated from the same distribution from which the normalized read counts for the major allele *M* was observed. Thus, the p-value 𝒫 to reject the null hypothesis was defined as:

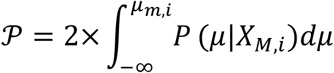

Based on Bayes rule, *P*(*μ*|*X_M,i_*) ∝ *P*(*X_M,i_*|*μ*)*P*(*u*). Assume the expected variance for the allele *M* is 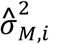 which is estimated as described in the last section. By combining the empirical probability *p*(*μ*|*X* = *X_M,i_*) with the distributions of *X* and *μ*, the empirical probability can be rewritten as a Gaussian distribution such that 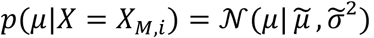 where the mean 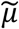 is:

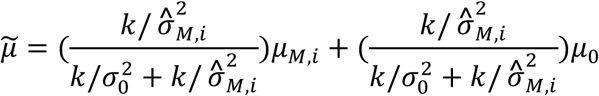

where *k* is the number of eCLIP replicates, and the variance 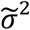 is:

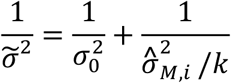

Since the prior probability of *μ* is unknown, we assumed that 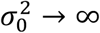 Thus, the posterior probability to observe the mean *μ_m,i_* for the minor allele *m* given *X_M,i_* is:

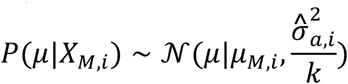

To adjust for multiple testing, FDRs were calculated. A minimum FDR of 10% was required to call significant ASB events in the ENCODE data sets.

### Identification of heterozygous SNVs in eCLIP reads

BEAPR includes a procedure to call heterozygous SNVs directly from the eCLIP data. As candidate SNVs, we obtained known SNPs or mutations from various databases including dbSNP, GTEx, TCGA, ExAC, and COSMIC. For each cell line, we pooled all the SMInput data sets together and calculated the allelic read counts at all candidate SNV locations. A candidate SNV was predicted as a heterozygous SNV in the cell line if the total read coverage was at least 10 and the allelic ratio of the reference allele was between 0.25 and 0.75. In addition, we used whole-genome DNA sequencing data of the two cell lines to identify heterozygous SNVs using the same method as in our previous work^14^.

### Simulation of allele-specific read counts

To simulate allele-specific read counts that mimic those in actual eCLIP data sets, we used the eCLIP data of SRSF1 in the K562 cell line generated by ENCODE. eCLIP peaks were identified as described above. For heterozygous SNPs (dbSNP 144) located in the peaks, we obtained their total read coverage in each replicate. The empirical total read coverage distributions were used to generate independent sets of simulated read counts. For each simulated SNP, its total read coverage was sampled from the above distribution. Its allelic ratio was set to be 0.5, unless it is a simulated ASB SNP (with allelic ratio being 0.7, 0.8 or 0.9). The allelic read counts for each SNP were determined using a zero-truncated negative binomial distribution with the expected variance set to be equivalent to the observed variance between the two replicates of SRSF1 eCLIP as a function of total read coverage.

### Evaluation of performance

Using simulated read counts, the overall performance of ASB prediction was evaluated in terms of precision (*PRE*), *PRE* = *TP/*(*TP* + *FP*), and recall (*REC*), *REC* = *TP/*(*TP* + *FN*), where *TP* is the number of true positives, *FP* is the number of false positives and *FN* is the number of false negatives. The area under the precision-recall curve (AUC) was calculated. We used this AUC value instead of that of a receiver operating characteristic (ROC) curve, because the numbers of SNVs with ASB or not were extremely unbalanced. A recent study suggested that precision-recall curves were more informative than ROC curves on unbalanced data^33^. Moreover, we also evaluated the prediction methods by sensitivity (SEN), *TP/*(*TP* + *FN*), and specificity (SPE), *FP/*(*TN* + *FP*), where *TN* is the number of true negatives. We reported sensitivity at 90% specificity (SEN90) and specificity at 90% sensitivity (SPE90) as additional performance metrics to assess the methods.

### Motif analysis

The position-specific motif enrichment plots (Fig. 3) were generated as follows. Within each peak harboring an ASB event, k-mer occurrence at each position flanking the ASB SNV was counted. Note that at the ASB SNV, the sequence of the major allele in eCLIP was used. The k-mers used here are pentamers identified by RBNS for each RBP^9^. The frequency of the RBNS pentamers in all ASB regions of an RBP was calculated. As controls, we randomly picked a genomic sequence to match each region with ASB in terms of the type of region (e.g., intron, 3’ UTR etc) and GC content (±10%). A total of 10 random sets of sequences were selected, with each set containing the same number of sequences as the number of ASB regions of an RBP. The fold enrichment of RBNS pentamers in ASB regions relative to the random controls was calculated and visualized in Fig. 3.

### Minigene reporters for splicing assays

For exon skipping events, the candidate exon and ~400nt upstream and downstream flanking introns were amplified using HeLa or K562 genomic DNA. After double digestion by HindIII and SacII or EcoRI and SacII, the DNA fragments were sub-cloned into pZW1 splicing reporter plasmids^34^. For intron retention events, the candidate intron and its flanking exons were cloned into the pcDNA3.1 plasmids. Final constructs were sequenced to ensure that a pair of plasmids containing the two alternative alleles of the SNV was obtained.

### Transfection, RNA extraction, Reverse transcription, and PCR

Minigene constructs were transfected into >90% confluence HeLa cells using Lipofectamine 3000 (ThermoFisher Scientific, L300015). Cells were harvested 24 h post transfection and total RNA was isolated using TRIzol (ThermoFisher Scientific, 15596018) followed by Direct-zol RNA Mini prep (Zymo Research, R2072). cDNA was prepared from 2 μg of total RNA by SuperScript IV First-Strand Synthesis System (ThermoFisher Scientific, 18091050) and one twentieth of the cDNA was used as template to amplify both inclusion and exclusion of the candidate exon by PCR of 28 cycles.

### Gel electrophoresis and quantification

Five microliter of PCR product was loaded onto 5% polyacrylamide gel and electrophoresis at 70 volt for one and a half hours. The gel was then stained with SYBR^®^ Safe DNA Gel Stain (ThermoFisher Scientific, S33102) for 30 min before imaging via Syngene SYBRsafe program (Syngene). Expression level of spliced isoforms was estimated using the ImageJ software (http://imagej.nih.gov/ij/). Inclusion or intron retention rate (% inclusion) of the target exon was calculated as the intensity ratio of upper/(upper+lower) bands.

### Bi-directional reporter constructs for 3’UTR analysis

To test the function of ASB events in 3’ UTRs, approximately 700-1000nt of the 3’ UTR regions including the ASB SNV were amplified using genomic DNA extracted from HMLE, HeLa or K562 cells. Site mutations were generated for alternative alleles for each SNV using overlap-extension PCR. After double digestion by ClaI and SalI-HF, the DNA fragments were sub-cloned into the 3’UTR of mCherry in the bi-directional reporter plasmid pTRE-BI-red/yellow that encodes for both mCherry and eYFP^35^. Final constructs were sequenced to ensure that a pair of plasmids containing the two alternative alleles of the SNV was obtained.

### Real-time PCR

The real-time PCR reaction was performed using SsoAdvanced Universal SYBR Green Supermix (Bio-Rad, 172-5270) and CFX96 Touch Real-Time PCR detection system (Bio-Rad) according to manufacturer’s instructions. The mRNA expression level associated with each allele of the ASB SNV was measured by the mCherry expression levels, which was normalized against that of eYFP.

### Purification of recombinant human PTBP1

The human PTBP1-pET28a expression vector was a gift from Dr. Douglas Black. It was transformed into BL21 Star (DE3) competent cells (ThermoFisher Scientific, C602003). Protein induction was carried out via 1mM IPTG treatment in 50mL cultured cells (O.D = 0.8) for 16 h at 215 rpm at 28°C. Next, cultured cells were centrifuged at 7000 × g for 5 min at 4°C and the pellets were resuspended with ice-cold 5mL lysis buffer (1 × BugBuster, 20mM Sodium Phosphate pH 7.7, 500mM NaCl, 20mM Imidazole, 1mM DTT, 0.5 × protease inhibitor cocktail, 100μg/mL lysozyme, 100U DNAse I). After 30 min incubation, the lysate was disrupted using three times sonication at 30% amplitude for 30 sec with 1 sec pulse. Subsequently, the lysate was centrifuged at 15,000 × g for 15 min at 4°C. The supernatant was collected and filtered using 0.45μm syringe filter. The sample was loaded into the HisTrap HP column (GE Healthcare, 17-5247-01) using Biologic LP system (Bio-Rad, 7318304) and washed with 20 mL buffer A (20mM Sodium Phosphate pH 7.7, 500mM NaCl, 20mM Imidazole, 1mM DTT). The sample was eluted with 500mM imidazole in buffer A. Purity of the recombinant PTBP1 protein was determined by SimplyBlue SafeStain (ThermoFisher Scientific, LC6060) and western blot using anti-HIS antibody (Santa Cruz Biotech, sc-8036, 1:500 dilution). Clean fractions (E28 and E33, Fig. S7) were combined (~3mL). Salt and small size of non-specific proteins were removed by incubating in 20K Slide-A-Lyzer dialysis cassette (ThermoFisher Scientific, 66003) with 1L Buffer A in a cold room overnight. Protein concentration was measured by Pierce Coomassie (Bradford) protein assay kit (ThermoFisher Scientific, 23200) and Turner spectrophotometer SP-830.

### In vitro transcription of PTBP1 target RNA

ASB candidates overlapping with PTBP1 binding motif were selected and 100uM of sense and antisense oligos including T7 promoter were annealed with oligo annealing buffer (10mM TRIS-HCl pH 8.0, 1mM EDTA pH 8.0, 100mM NaCl) at 95°C for 5min in a heat block then cooled slowly to 28°C for 2hr. In vitro transcription was performed using 1μg of annealed oligos and HiScribe T7 high yield RNA synthesis kit. In vitro synthesized RNAs were treated with 10U RNAse-free DNAse I (ThermoFisher Scientific, EN0525) at room temperature for 30min, then purified by RNA clean & concentrator-5 Kit (Zymo Research, R1015). Next, RNA samples were treated with 10U shrimp alkaline phosphatase (NEB, M0371S) at 37°C for 1 hr and then labeled with 0.4μl of gamma 32P-ATP (7000Ci/mmol, MP Biomedicals) using 20U T4 polynucleotide kinase (NEB, M0201S). Subsequently, RNA probes were purified using 5% Urea PAGE extraction and RNA clean & concentrator-5 Kit. RNA concentration was measured by Qubit 2.0 fluorometer (ThermoFisher Scientific).

### Electrophoretic Mobility Shift Assay (EMSA)

The purified RNA probes (20pmol) and recombinant PTBP1 protein (0, 0.6, 1.2, 2.5, and 5 μg) were incubated in 15μl of buffer A (20mM Sodium Phosphate pH 7.7, 500mM NaCl, 20mM Imidazole, 1mM DTT, 0.1× protease inhibitor cocktail, 10U RNAse inhibitor) at 28°C for 30 min, then loaded onto 5% TBE-PAGE and ran at 75V for 1.5h. The gel was processed without drying, covered with clear folder and exposed to X-ray film at ‒80°C.

## Acknowledgements

We thank members of the Xiao, Graveley and Yeo laboratories for helpful discussions and comments on this work. This work was funded by the National Human Genome Research Institute ENCODE Project, grants U01HG009417 to X.X. and U54HG007005 to B.R.G. and G.W.Y. X.X. was partially supported by grants from the NIH (R01HG006264, R01AG056476). E.L.V.N. is a Merck Fellow of the Damon Runyon Cancer Research Foundation (DRG-2172-13) and is supported by a K99 grant from the NIH (HG009530). G.A.P. is supported by the NSF graduate research fellowship.

## Data availability

All data sets used in this study can be obtained from the ENCODE project website at http://www.encodeproject.org.

